# C3G dynamically associates with Nuclear speckles and regulates mRNA splicing

**DOI:** 10.1101/159269

**Authors:** Dhruv Kumar Shakyawar, Bhattiprolu Muralikrishna, Vegesna Radha

## Abstract

C3G (RapGEF1), essential for mammalian embryonic development, is ubiquitously expressed and undergoes regulated nucleo-cytoplasmic exchange. Here we show that C3G localizes to SC35 positive nuclear speckles, and regulates splicing activity. Reversible association of C3G with speckles was seen upon inhibition of transcription and splicing. C3G shows partial colocalization with SC35, and is recruited to a chromatin and RNase sensitive fraction of speckles. Its presence in speckles is dependent on intact cellular actin cytoskeleton, and is lost upon expression of the kinase, Clk1. Rap1, a substrate of C3G, is also present in nuclear speckles and inactivation of Rap signalling by expression of GFP- Rap1GAP, alters speckle morphology and number. Enhanced association of C3G with speckles is seen upon GSK3β inhibition, or differentiation of C2C12 cells to myotubes. CRISPR/Cas9 mediated knockdown of C3G resulted in decreased splicing activity and reduced staining for SC35 in speckles. C3G knockout clones of C2C12 as well as MDA-MB- 231 showed reduced protein levels of several splicing factors compared to control cells. Our results identify C3G and Rap1 as novel components of nuclear speckles and a role for C3G in regulating cellular RNA splicing activity.

**Summary:** Nuclear speckles are sites for pre-mRNA splicing. We provide evidence for localization and function of a Ras family GTPase, Rap1 and its exchange factor C3G in nuclear speckles.

## Introduction

Many molecules function in signalling pathways through dynamic nucleo-cytoplasmic exchange to regulate a variety of nuclear functions like chromatin organization, gene expression and RNA processing. Within the nucleus, proteins may be present in the nucleoplasm, or associated with nuclear sub-structures like chromatin, nuclear matrix, nuclear membrane, nucleoli, Cajal bodies, or nuclear speckles (Handwerger and Gall, 2006). Their localization often provides insights into the functions they perform in the nucleus. Processes like replication, transcription and DNA repair take place in distinct nuclear regions, and are generally defined by dynamics of chromatin remodelling and nuclear architecture (Stein et al., 2003).

Nuclear speckles are defined, irregularly shaped structures present in the inter-chromatin regions (Spector and Lamond, 2011). They are complexes of a large number of proteins and snRNA, and are sites of pre-mRNA splicing in cells (Spector, 1993; Zhou et al., 2002). Precise excision of introns from the primary transcript is important to generate spliced translatable mRNAs in eukaryotes. RNA splicing is a regulated process mediated by a multiprotein complex called spliceosome which facilitates the chemical removal of introns (Misteli et al., 1997). Two classes of proteins, the SR family, and hnRNPs, primary localize to speckles and engage in splicing activity. In addition, several other regulatory molecules like kinases and phosphatases also localize to speckles (Spector and Lamond, 2011). Some of these proteins have signature sequences that enable their targeting to speckles (Eilbracht and Schmidt-Zachmann, 2001; Jagiello et al., 2000; Salichs et al., 2009), and phosphorylation plays a major role in regulating localization to speckles (Mermoud et al., 1994; Stamm, 2008). While some proteins are constitutive to speckles, and play a role in functional mRNA biogenesis, others show restricted expression and tissue specific alternate splicing events (Castle et al., 2008; Wang et al., 2008). Speckle size and structure are dependent on transcription and splicing activity in cells, and active transcription takes place in the periphery of speckles (Lamond and Spector, 2003; Misteli et al., 1997). RNA transcription and splicing are altered upon transient heat stress, which also affects the localization of various factors to speckles (Biamonti and Caceres, 2009; Lallena and Correas, 1997; Spector et al., 1991; Velichko et al., 2013).

C3G is a ubiquitously expressed guanine nucleotide exchange factor (GEF) that regulates important cellular functions like cytoskeletal remodelling, adhesion, cell proliferation and apoptosis (Radha et al., 2011). It is a 140 kDa protein with a catalytic domain at the C-terminus, and a central protein interaction domain encompassing multiple poly-proline rich tracts. The N-terminus has an E-cadherin binding domain, and the non-catalytic domains serve to negatively regulate its catalytic activity (Hogan et al., 2004; Ichiba et al., 1997). In addition, C3G activity is regulated by tyrosine phosphorylation and membrane localization (Ichiba et al., 1999; Ichiba et al., 1997). Functions of C3G are dependent on its catalytic and/or protein interaction properties. C3G targets GTPases Rap1, Rap2, R-Ras, and TC10 (Gotoh et al., 1995; Gotoh et al., 1997; Mochizuki et al., 2000); and interacts with signalling molecules like Hck, Src, Crk, c-Abl, TC-48, and β-Catenin (Dayma et al., 2012; Knudsen et al., 1994; Mitra et al., 2011; Radha et al., 2007; Shivakrupa et al., 2003; Tanaka et al., 1994). We showed that C3G interacts with, and suppresses β-catenin activity, a determinant of cell fate decisions (Dayma et al., 2012). C3G levels increase, and it is required for differentiation of muscle and neuronal cells (Radha et al., 2008; Sasi Kumar et al., 2015). C3G is essential for early embryonic development in mammals, as mice lacking C3G do not survive beyond 6-7.5dpc (Ohba et al., 2001).

Indirect immunofluorescence experiments have shown that cellular, as well as over-expressed C3G localizes primarily to the cytoplasm (Hogan et al., 2004; Radha et al., 2004). The primary sequence of C3G has functional NLSs, and, fractionation of a variety of cells has shown the presence of C3G in the nuclear compartment (Shakyawar et al., 2017). Inhibition of CRM1 dependent export as well as GSK3β activity, caused an increase in nuclear levels of C3G. In an attempt to understand functions for C3G in the nucleus, we examined its localization to distinct sub-nuclear structures. Our findings demonstrate that C3G and its target Rap1 show dynamic association with nuclear speckles dependent on cellular transcription activity. Differentiation of myocytes to myotubes enhances association of C3G with nuclear speckles and knockdown of C3G causes reduction in cellular splicing activity.

## Results

### Inhibition of nuclear export causes enrichment of C3G in SC35 positive speckles

The dynamic exchange of C3G between nuclear and cytoplasmic compartments prompted us to study the localization of C3G to distinct sub-nuclear domains, to understand its function in the nucleus. Since C3G localized predominantly to inter chromatin regions upon inhibition of nuclear export, we examined its localization to nuclear speckles by co-staining with antibody for SC35(SRSF2), a spliceosomal protein of SR family, and an established nuclear speckle marker. This antibody recognizes a phospho-epitope of SC35 (Fu and Tom, 1990). A commercial antibody raised against N-terminal region of C3G that specifically recognizes endogenous C3G, was used (Fig. 1A). This antibody recognizes endogenous C3G in methanol fixed cells when examined by indirect immunofluorescence. Treatment of MDA-MB-231 cells with nuclear export inhibitor Leptomycin B (LMB) showed prominent staining for C3G in nuclear foci positive for SC35, upon analysis by confocal microscope (Fig. 1B). This pattern was seen in a large number (50-60%) of cells. Untreated cells show cytoplasmic and weak nuclear staining. The efficacy of LMB treatment was validated by observing nuclear localization of NF-κB a protein regulated by CRM1 dependent nuclear export (Fig. S1A). Localization of C3G to speckles in response to LMB, was also seen in other human cell types like MCF-7 (Fig. S2A). Co-localization of endogenous C3G was also seen with overexpressed SC35-GFP upon LMB treatment (Fig. S2B). All cells expressing SC35-GFP showed localization of C3G to splicing factor compartments.

**Fig. 1.**
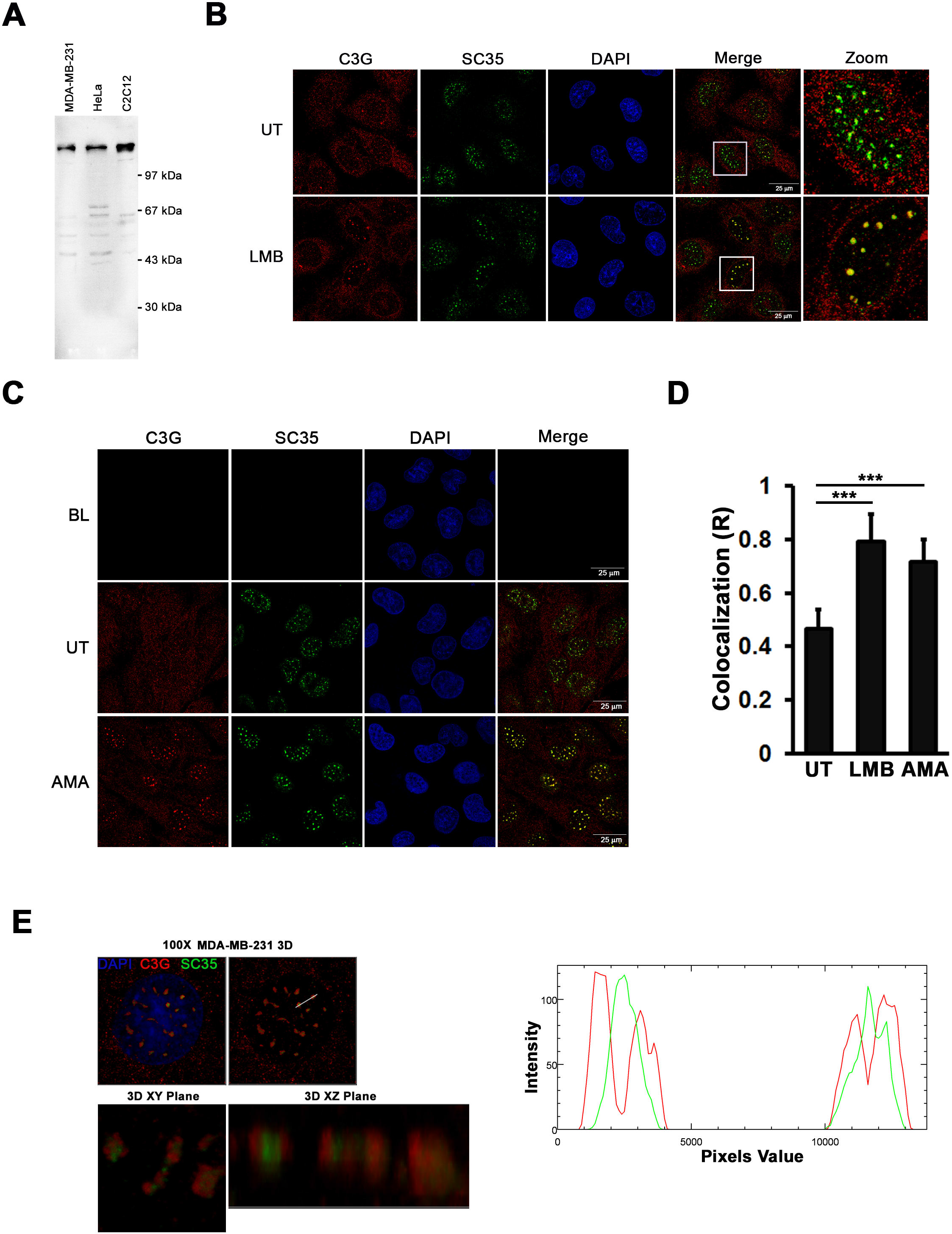
C3G localizes to SC35 nuclear speckles. **A.** MDA-MB-231, HeLa, and C2C12 cell lysates were subjected to immunoblotting and probed for C3G expression using C3G antibody. **B**. Normally growing MDA-MB-231 cells (UT) on coverslips or those treated with LMB, were fixed and immunostained for C3G and SC35. Panels show optical sections taken using a confocal microscope. **C**. MDA-MB-231 cells were subjected to α-Amanitin treatment; and immunostained for C3G and SC35. BL indicates cells processed similarly without addition of primary antibodies. **D**. Bar Diagram represents the Pearson correlation coefficient of colocalization between C3G and SC35 analysed from multiple images, upon indicated treatments. **E**. Optical sections (z-plane step, 0.30 µm) of nuclei from amanitin treated cells were captured on Leica SP8 confocal microscope, and were reconstructed to form a three-dimensional image. Three-dimensional visualization of speckle regions of cells dually labelled with antibodies against C3G (Red) and SC35 (Green) in XY or XZ plane are shown. Line scans showing local intensity distributions of C3G in red and SC35 in green in the ROI drawn across two speckles are shown to the right of the panels.

### Inhibition of transcription results in enhanced localization of C3G to speckles

The shape & size of speckles is known to change depending on cellular transcription levels (Melčák et al., 2000). Inhibition of transcription causes formation of large intra nuclear foci containing splicing factors in cells (Carmo-Fonseca et al., 1992; Hall et al., 2006; Spector, 1993). This is a characteristic feature of nuclear speckles (IGC-inter chromatin granule clusters) which undergo reorganisation and fuse to form large structures which are more rounded and fewer in number. To examine if C3G localization to nuclear speckles responded to inhibition of transcription, α-Amanitin treated MDA-MB cells were dual stained with C3G and SC35antibodies. We observed a redistribution of C3G into enlarged speckles in response to inhibition of transcription, similar to that seen for SC35 (Fig. 1C). No signal was seen in speckles when C3G primary antibody was omitted, and cells were exposed to only secondary antibodies. Quantitation of extent of co-localization between C3G and SC35 showed a significant increase in LMB and amanitin treated cells (Fig. 1D). Inhibition of transcription by amanitin was confirmed by examining RNA pol II using the antibody, H5 which recognises the active Ser2phosphorylated form of RNA Pol II, (Fig. S1B) (Warren et al., 1992). Unlike under condition of inhibition of nuclear export, treatment with amanitin resulted in all the cells showing C3G in speckles. Localization of C3G to nuclear speckles was not specific to a particular cell type, as it was also observed in HeLa, and C2C12 cells (Fig. S3A-B). Endogenous C3G also showed colocalization in nuclear speckles with other ribo-nucleoproteins like U1 snRNP70, U2 snRNP B" and Sm snRNP (Y12) upon inhibition of transcription (Fig. S4A-D).

When examined at higher magnification, C3G was distributed irregularly in speckles and appeared to be present only partially co-localized with SC35. Individual speckles were examined by capturing optical z sections at 0.30 microns. 3D reconstruction showed distribution of C3G non-uniformly in the speckle, being prominently seen in the periphery covering the core region where SC35 is primarily located. This was also evident from XY & XZ projections (Fig. 1E, left panel). Quantitative assessment of fluorophore localization using a region of interest (ROI) was carried out to assess the spatial distribution of C3G and SC35 within speckles which showed that intensity peaks do not correspond though the signals overlapped (Fig. 1E). This data indicated that C3G localizes to discrete regions that show only partial overlap with SC35.

The dynamics of localization of C3G to speckles was examined using a reversible transcriptional inhibitor DRB (Sehgal et al., 1976). C3G showed reversible association, similar to SC35, as its prominence in speckles was reduced 1 hr after removal of DRB (DRB Rec) (Fig. 2A). Since C3G dynamically exchanges between nuclear & cytoplasmic compartments, we examined if the prominent association with speckles in response to α-amanitin treatment was due to enhanced nuclear translocation or retention. Levels of C3G in nuclear & cytoplasmic fractions were compared in untreated (UT) and amanitin treated cells. Unlike LMB treatment (Shakyawar et al., 2017), amanitin does not increase nuclear C3G levels, indicating that enhanced association with speckles was due to redistribution of nuclear C3G (Fig. 2B).

**Fig. 2.**
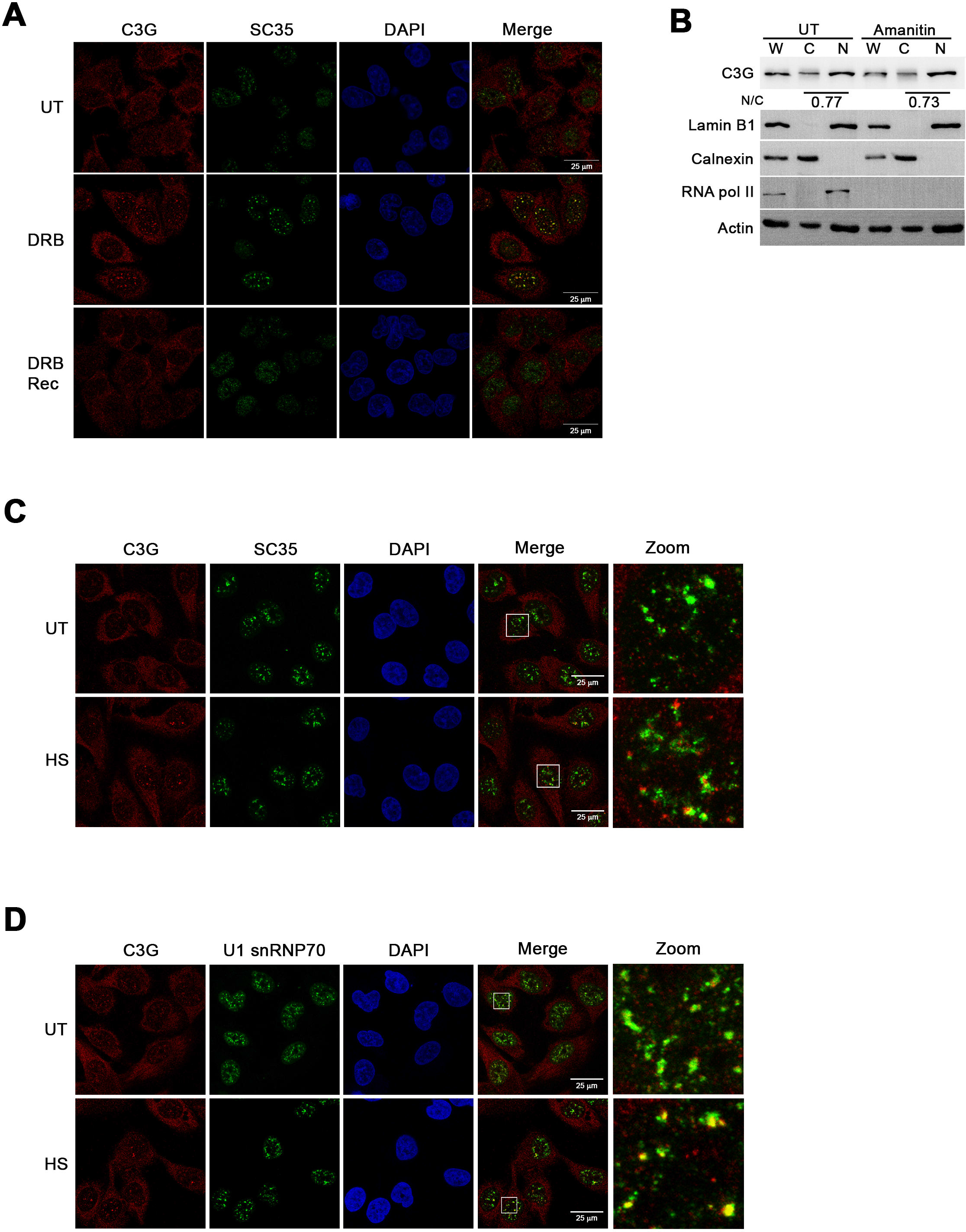
C3G shows reversible association with nuclear speckles. **A**. Untreated MDA-MB- 231 cells (UT) plated on coverslips were treated with DRB and either fixed immediately or after recovery from drug treatment for 1 hr (DRB Rec). Cells were subjected to indirect immunofluorescence to detect C3G and SC35 expression. **B.** MDA-MB-231 cells grown in the presence or absence of amanitin were subjected to cell fractionation. Lysates were processed for immunoblotting and probed for expression of indicated proteins. Calnexin and Lamin B1 were used to indicate the purity of the cytoplasmic and nuclear fractions respectively. H5 antibody (RNA pol II pS2) was used to show efficacy of α-amanitin treatment. Number indicate ratio of C3G levels in nuclear vs cytoplasmic fraction determined, based on protein loading. Effect of heat shock on C3G localization. MDA-MB- 231 cells plated on coverslips were grown at 37°C (UT) or subjected to heat shock (HS). Panel shows confocal images of cells immunostained with antibodies against C3G and SC35 **(C)** or C3G and U1 snRNP70 **(D)**.

SC35 and snRNPs respond differently to heat shock (HS), which is also known to inhibit transcription (Lallena and Correas, 1997; Spector et al., 1991). We examined the localization of C3G in response to heat shock, and observed that about 40-50% of cells showed the presence of C3G in alternate nuclear structures which were irregular and very few in number per cell. These structures did not colocalize with SC35 positive speckles (Fig. 2C). We also examined the colocalization of C3G and U1 snRNP70 under similar condition and found that unlike SC35, U1 snRNP70 partially colocalized with C3G in these alternate nuclear structures.

The rounding up of speckles and clustering of splicing factors upon inhibition of transcription is caused by their accumulation at storage sites, due to reduction in pre-mRNA levels. Reduction in splicing activity also results in similar changes in speckle morphology and number (O’keefe et al., 1994). We examined localization of C3G in cells treated with Isoginkgetin (IGK), a pre-spliceosome complex inhibitor (O’Brien et al., 2008). Reversible association of C3G with speckles was seen upon treatment of cells with IGK (Fig. 3A), suggesting a role for C3G in splicing. Formation of nuclear speckles is dependent on phosphorylation of proteins by the kinase Clk1, and exogenous expression of Clk1 causes redistribution of SR proteins out of speckles (Colwill et al., 1996). GFP-Clk1 and its catalytic mutant (GFP-mClk1) were expressed in MDA-MB cells, and examined for C3G expression in normal and amanitin treated cells. Just as described for other SR/SF proteins, expression of Clk1, but not its kinase inactive variant resulted in loss of speckle pattern of staining for C3G in amanitin treated cells (Fig. 3B). These results suggested that localization of C3G to the speckles is dependent on activity of Clk1 and may be coupled to distribution of SR factors.

**Fig. 3.**
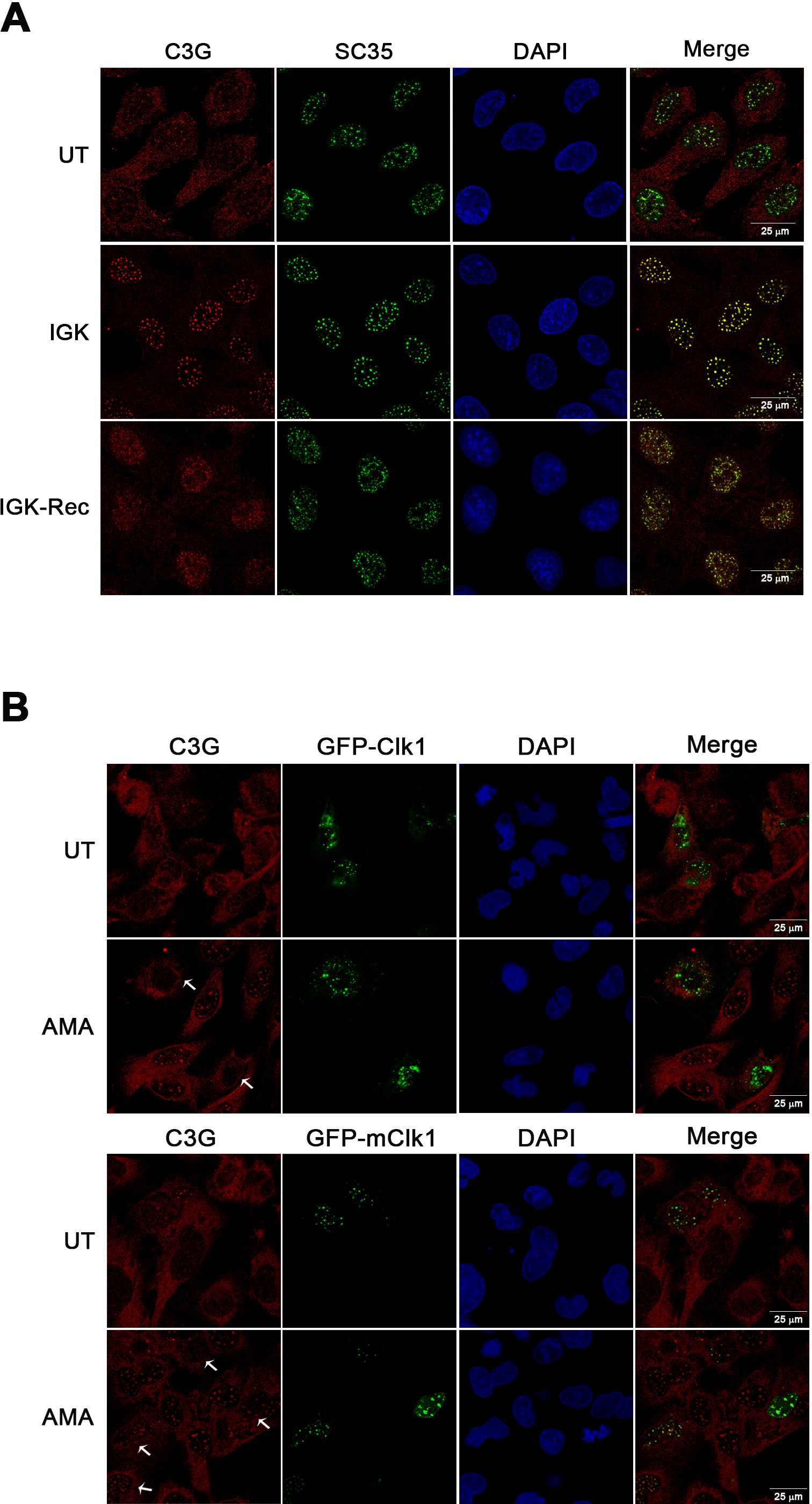
C3G localization to speckles increases upon inhibition of splicing and is lost upon overexpression of Clk1. **A**. MDA-MB-231 cells were either left untreated (UT) or treated with IGK. Cells were also allowed to recover for 6 h post drug washout (IGK-Rec). Cells were fixed and subjected to immunofluorescence with antibodies against C3G and SC35. Images of a single Z-section through the center of the nucleus acquired on a confocal microscope are shown in the panels. **B.** MDA-MB-231 cells expressing GFP-Clk1 or GFP- mClk1 construct were left untreated, or subjected to α-Amanitin treatment, fixed, and immunostained with antibody against C3G. Panels show confocal images of cells expressing C3G and transfected GFP tagged constructs. Arrows indicate GFP expressing cells in the C3G panels.

### C3G is associated with nuclear speckles dependent on intact chromatin and RNA

Molecules generally localize to speckles through association with proteins or RNA (Dye and Patton, 2001; Lallena and Correas, 1997). The association of C3G with speckles was examined by carrying out detergent extraction as well as sensitivity to DNase I & RNase A treatments. When α-Amanitin treated MDA-MB cells were extracted with detergent prior to fixation, C3G as well as SC35 were intact. Treatment with DNase I also did not alter C3G and SC35 staining (data not shown). When treatment with DNase I was followed by 0.4 M NaCl, the SC35 positive intra nuclear foci were intact but C3G staining was greatly reduced as some part of chromatin was lost from the nucleus (Fig. 4A). When digestion was followed by 2M NaCl treatment which removes all chromatin and retains only nuclear matrix, C3G was totally lost from speckles. SC35 foci were intact, indicating tight association with nuclear matrix. Removal of chromatin was confirmed by the loss of DAPI staining.

**Fig. 4.**
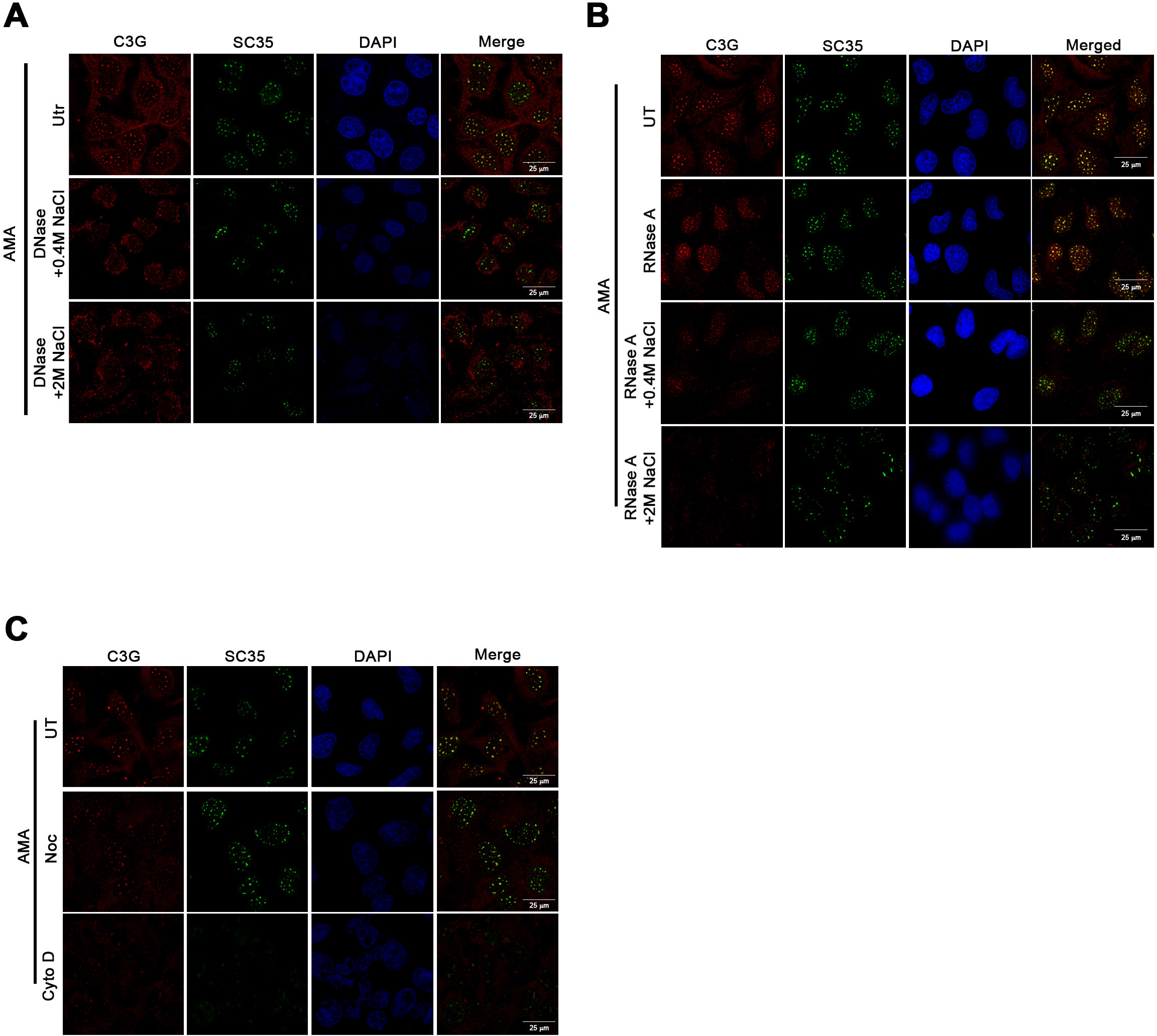
C3G localization to nuclear speckles is dependent on the presence of intact chromatin and RNA. Amanitin treated MDA-MB-231 cells were left untreated or subjected to DNAse **(A)** or RNAse **(B)** treatment followed by NaCl extraction, fixed, and immunostained for C3G and SC35. **C.** MDA-MB-231 cells were treated with α-Amanitin, and also exposed to Nocodazole or Cytochalasin D for 4 h prior to fixation. Immunofluorescence was carried out to detect C3G and SC35.

Upon RNase A treatment, speckle localization of C3G was reduced showing diffused nuclear staining, though most of the RNA was lost from the cells (Fig. 4B & S1C). When RNase treatment was followed by 0.4 M and 2M NaCl extraction, the enlarged intra nuclear foci formed by C3G were totally reduced, whereas SC35 foci were nearly intact (Fig. 4B). High salt extraction resulted in retention of SC35 staining only in sharp speckles. The efficacy of RNase A digestion was confirmed by the absence of staining with anti m3G antibody (Fig. S1C) which labels capped snRNAs. These results indicated that localization of C3G to speckles was dependent on the presence of intact chromatin and RNA in cells.

We examined the requirement of intact cellular microtubules and microfilaments for localization of C3G to speckles upon amanitin treatment. C3G staining in intranuclear large foci formed after α-amanitin treatment was partially reduced when MDA-MB cells were treated with Nocodazole, an inhibitor of microtubule polymerisation. There was no effect on SC35 foci. When treated with Cytochalasin D (an inhibitor of Actin polymerisation), C3G as well as SC35 showed a weak diffuse pattern, without prominent localization to speckles (Fig. 4B). The efficacy of Cytochalasin D and Nocodazole treatments was verified by staining similarly treated parallel coverslips for F-actin and α-tubulin respectively (data not shown).

### Rap1, a target of C3G localizes to speckles

Since C3G is a GEF for Ras family GTPases, we examined if Rap1, a target of C3G localizes to nuclear speckles. Localization of Rap1 to the nucleus has been shown earlier (Lafuente et al., 2007). Endogenous Rap1 showed non-uniform staining throughout the cell when methanol fixed cells were examined by immunofluorescence, similar to that shown earlier (Bivona et al., 2004), but in amanitin treated cells, prominent localization was seen at nuclear speckles (Fig. 5A). To determine if activated form of Rap1 is present in speckles, we examined localization of activated (GTP bound) Rap1 by expressing a GFP fusion protein of Ral-GDS-RBD, which binds specifically to activated molecules of Rap1 in cells, and can be observed as enhanced GFP signals (Bivona TG, 2004). We observed prominent GFP fluorescence at nuclear speckles, specifically in α-amanitin treated cells (Fig. 5B), in addition to diffuse signals seen throughout the cell. In untreated cells, intense GFP signals were seen in punctate cytoplasmic structures, which are likely to be endosomes/multi-vesicular bodies as described earlier (Pizon et al., 1994). These results suggest a function for C3G and its target, Rap1 in nuclear speckles.

**Fig. 5.**
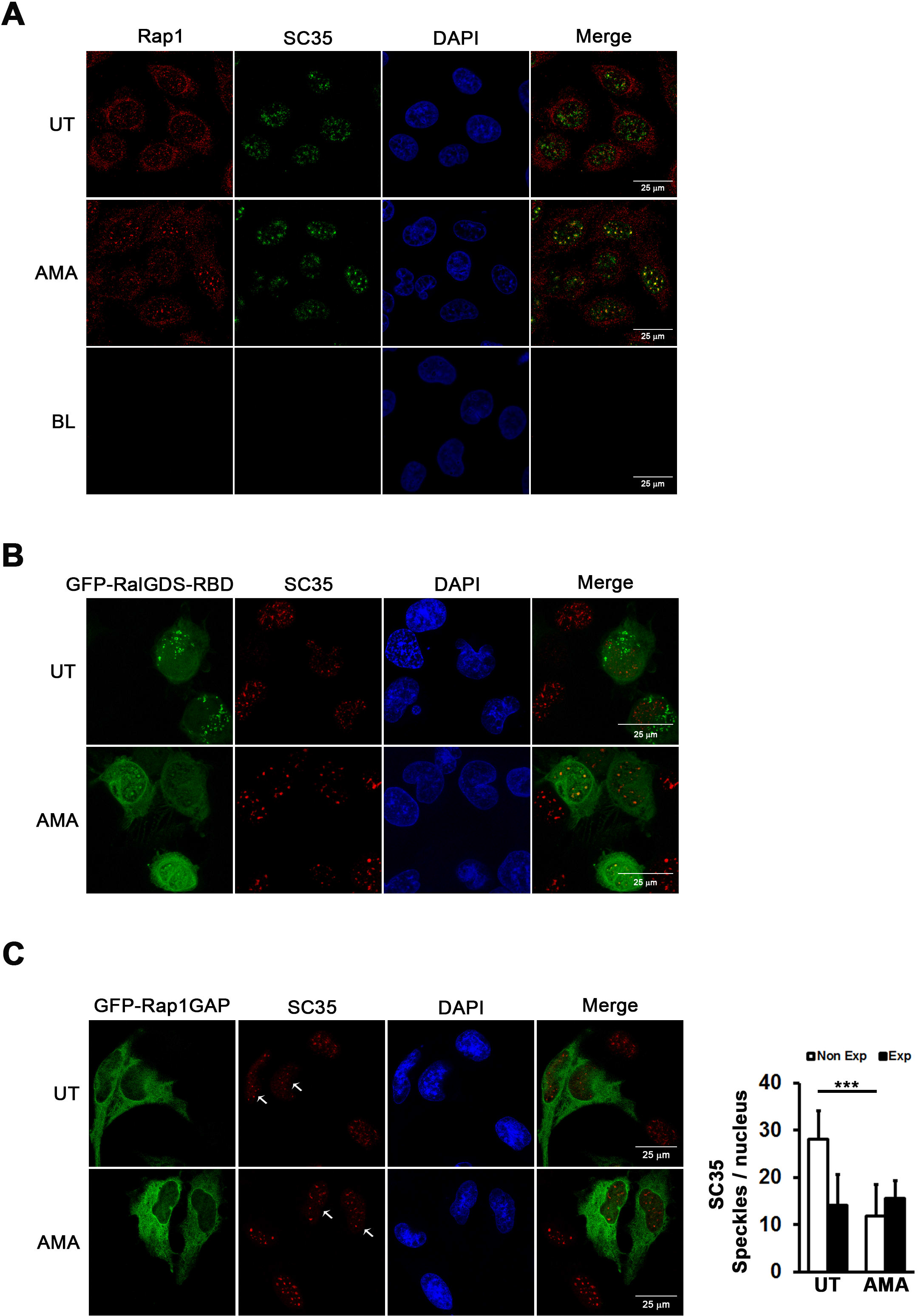
Rap1 a target of C3G localizes to nuclear speckles. **A**. MDA-MB-231 cells were treated with (AMA) or left untreated (UT) and immunostained for Rap1 and SC35. BL indicates secondary antibody control where cells were processed without addition of primary antibodies. **B**. GFP-RalGDS-RBD transfected MDA-MB-231 cells were treated with or without α-Amanitin, fixed with formaldehyde and immunostained with SC35. **C.** MDA-MB- 231 cells transfected with GFP-Rap1GAP were treated with or without amanitin, fixed with methanol and immunostained for expression of SC35. Arrows in SC35 panel show GFP- Rap1GAP expressing cells. Bar diagram shows quantitation of number of speckles per nucleus in expressing and non-expressing cells using data obtained from large number of cells from three independent experiments. ***p < 0.001.

This was further validated by examining SC35 speckles in cells expressing GFP-Rap1GAP, a protein known to inhibit Rap activation dependent downstream signalling. We compared structure and number of SC35 speckles in MDA-MB cells expressing GFP-Rap1GAP, (in normally growing and under conditions of transcription inhibition) with those that do not express GFP-Rap1GAP. As shown in Fig. 5C, GFP-Rap1GAP expressing cells show more compacted and significantly fewer speckles compared to non-expressing cells. Difference in speckle morphology and number were not seen under conditions of amanitin treatment, with both expressing and non-expressing cells showing fewer and more rounded speckles.

### C3G is required for pre-mRNA splicing

Earlier work from our laboratory showed that C3G translocates to the nucleus in response to LiCl, an inhibitor of GSK3β (Shakyawar et al., 2017). Upon LiCl treatment which causes increased expression of many genes, we observed that C3G shows enhanced localization to speckles (Fig. 6A). SC35 localization was also altered upon LiCl treatment as shown earlier (Hernández et al., 2004). Efficacy of LiCl treatment was tested by immunostaining of β-catenin, which shows nuclear translocation in response to LiCl treatment (Data not shown).

**Fig. 6.**
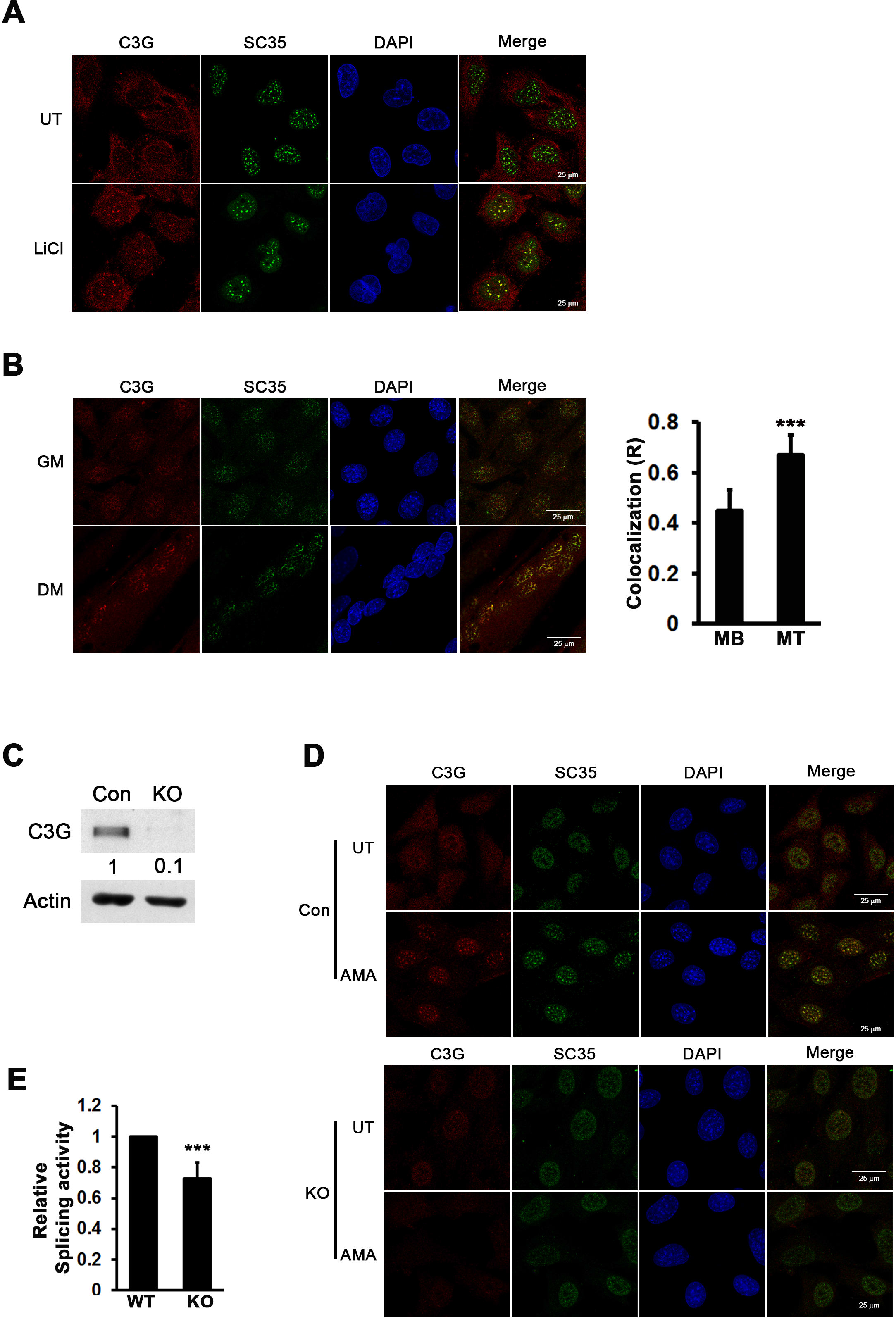
C3G associates with nuclear speckles upon myocyte differentiation and is required for splicing activity. **A.** MDA-MB-231 cells grown on glass coverslips were subjected to LiCl treatment prior to fixation, and immunostained for expression of C3G and SC35. Images show confocal sections of immunostained cells. **B.** C2C12 cells were grown in the presence of growth medium (GM) or differentiation medium (DM) for 72 h, fixed and immunostained for expression of C3G and SC35. The colocalization coefficient between C3G and SC35 in myoblasts and myotubes was quantitated using large number of cells from 3 independent experiments is shown in bar diagram. ***p < 0.001. **C.** Immunoblot of lysates from control (Con) and C3G KO clone (KO), probed for the expression of C3G; actin was used as loading control. Numbers indicate relative amount of C3G in KO cells compared to control cells. **D**. SC35 staining is altered in C3G KO cells. Control and C3G KO clones of C2C12 were treated with or without α-Amanitin, fixed and immunostained to detect C3G and SC35. Panels show images acquired on a confocal microscope. **E.** Effect of C3G KO on cellular splicing activity. Control and C3G KO clones transiently transfected with pTN24 vector for 48 h and lysates were assayed for luciferase and β-gal activity. Bar diagram shows relative luciferase to β-gal ratio averaged from three independent experiments. ***p < 0.001.

Inhibition of GSK3β promotes myogenic differentiation and SC35 speckles show reorganization during myocyte differentiation, becoming more mis-shapen and larger (Homma et al., 2016; van der Velden et al., 2008). We have earlier shown that C3G is required for C2C12 differentiation, and that C3G translocates to nuclei upon differentiation of myocytes to form myotubes (Sasi Kumar et al., 2015). Co-localization studies showed that C3G localization in SC35 positive speckles of myotubes is higher compared to that seen in undifferentiated cells (Fig. 6B).

Splicing activity and structure of nuclear speckles changes during myocyte differentiation (Homma et al., 2016). Based on its dynamic localization to speckles, we hypothesized a role for C3G in mRNA processing. We examined the role of cellular C3G in regulating activity in nuclear speckles by using a clone of C2C12 cells where C3G was knocked down using CRISPR/Cas9 technology (Fig. 6C). Unlike control cells, these cells do not fuse to form myotubes when grown in differentiation medium (Shakyawar et al., 2017). Compared to normal C2C12, cells with reduced cellular C3G showed very poor staining for SC35 speckles (Fig. 6D). In amanitin treated KO cells also, SC35 speckles were weakly stained and less distinct throughout the nucleus compared to WT cells.

We went on to determine if loss of C3G also impacted cellular splicing activity using a double reporter splicing assay plasmid, pTN24. Ratio of the activities of the two reporter genes, luciferase and β-gal, gives information on the extent of splicing activity in cells. This plasmid was expressed in WT and C3G KO C2C12 cells and splicing efficiency determined. C3G KO cells showed significantly lower splicing activity, suggesting a role for C3G in regulating RNA splicing. (Fig. 6E).

The reduced levels of SC35 seen in C3G knockout cells could be either a consequence of inefficient assembly into speckles, or due to changes in expression level of SC35. We therefore examined total protein levels of SC35, and a few other splicing factors in WT C2C12 cells and C3G KO cells. As shown in Fig. 7A, expression of many of the tested splicing factors was significantly lower in cells lacking C3G. Similar reduction was also observed in C3G KO clones of MDA-MB-231 cells (generated using CRISPR/Cas9 technology) indicating that the effect was not restricted to a particular cell type (Fig. 7B). These results suggested that cellular C3G functions to regulate expression of splicing factors.

**Fig. 7.**
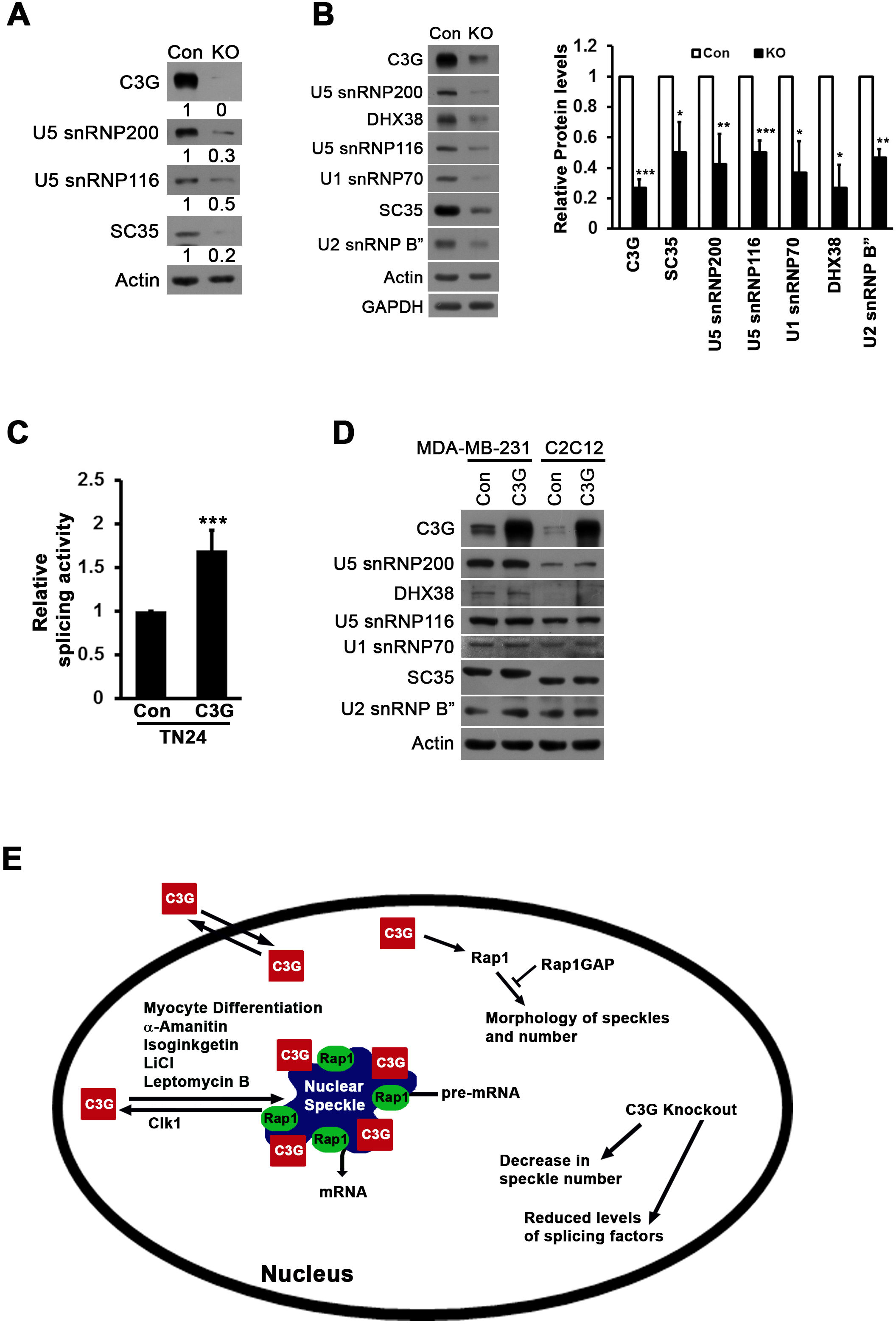
Altered expression of splicing factors in C3G knockout cells. **A.** Lysates of C2C12 control and C3G KO clones were subjected to Western blotting and probed for C3G, U5 snRNP200, U5 snRNP116 and SC35. Actin was used as loading control. Numbers indicate relative protein levels in KO compared to control. **B.** Whole cell lysates of MDA-MB-231 control and C3G KO clones, probed for the expression of the indicated proteins. Bar diagram shows relative levels of the proteins averaged from three independent experiments. *p < 0.05; **p < 0.01; ***p < 0.001. **C.** Effect of C3G overexpression on splicing factor levels and activity. HEK-293T cells were cotransfected with pTN24 along with C3G-Flag or control vector. After 48 h, lysates were assayed for luciferase and β-gal activity. **D.** MDA-MB-231 and C2C12 cells were transiently transfected with C3G-Flag construct for 30 h. Lysates were subjected to immunoblotting, and blot was probed for indicated proteins. **E.** Schematic figure depicting dynamic localization of C3G and its effector GTPase Rap1 to nuclear speckles. Inhibition of transcription and splicing as well as physiological stimuli enhance localization of C3G to SC35 positive speckles, whereas expression of active Clk1 disrupts amanitin induced C3G localization to speckles. Active form of Rap1 is present in nuclear speckles and inactivation of Rap1 mediated by expression of Rap1GAP resulted in altered morphology of speckles and significant decrease in number. Cells lacking C3G due to CRISPR/Cas9 mediated knockdown show reduced levels of splicing factors and decrease in splicing activity.

Presuming that over-expression of C3G would have some consequence on splicing factor levels, we examined levels of these proteins in lysates of MDA-MB-231 and C2C12 cells, overexpressing C3G. It was observed that none of the proteins showed any significant change in their cellular levels upon C3G overexpression (Fig. 7C). We also examined the effect of enhancing cellular levels of C3G on splicing activity and observed that co-expression of C3G with TN24 caused a significant increase in splicing activity (Fig. 7D).

## Discussion

Molecules that exchange between the nuclear & cytoplasmic compartments, regulate important processes in the nucleus. As a follow up of our study identifying nuclear localization of C3G in response to certain physiological stimuli, we carried out a detailed investigation into nuclear functions of C3G. In this study, we identify a novel function for C3G in the nucleus, where it localizes to SC35 positive speckles, and regulates splicing activity. Localization of C3G to speckles is a feature conserved in cells derived from various tissue types, as well as in murine and human cells.

Upon inhibition of nuclear export, C3G is retained in the nucleus, with prominent localization seen with SC35 speckles in several cells. Just as in the case of a variety of other molecules involved in RNA splicing activity (Carmo-Fonseca et al., 1992), C3G also shows enhanced localization to speckles upon inhibition of transcription or splicing. Cell fractionation indicated that inhibition of transcription did not particularly increase nuclear levels of C3G, suggesting that nuclear C3G exchanged between nucleoplasm & speckles. The reversibility of release from speckles seen upon removal of transcription or splicing activity inhibition, similar to other splicing factors suggests a common mechanism of dynamic exchange between speckles and nucleoplasm. The fact that C3G is not prominently seen in speckles of exponentially growing cells indicates that it is in constant flux between nucleoplasm and speckles.

In response to heat shock, transcriptional reprogramming occurs & general RNA transcription and splicing are also inhibited (Velichko et al., 2013). It therefore came as a surprise that upon heat shock, C3G did not localize to SC35 positive speckles and appears to be present in alternate granular structures in the nucleus which showed partial colocalization with U1 snRNP70. It is reported that few splicing factors but not SC35, re-localize to nuclear stress granules which are sites of HSF1 localization and activity upon heat shock (Chiodi et al., 2000; Denegri et al., 2001; Weighardt et al., 1999). It is suggested that splicing factors are sequestered in speckles, and made available in response to specific stimuli (Spector and Lamond, 2011). If C3G is moving to nuclear stress granules in response to heat shock, when enhanced transcription of chaperones takes place, it is tempting to speculate that C3G enables gene specific transcription in response to heat stress.

Though C3G localizes to speckles, it does not show superimposition with SC35, and is present in a spatially heterogenous fashion (more in peripheral regions, compared to SC35), suggestive of being involved in distinct functions within the speckle (Mintz and Spector, 2000). C3G is released from speckles upon extraction of chromatin, or RNA from nuclei, and differs from SC35 with respect to its extractability from speckles. Removal of C3G from speckles upon mild extraction conditions is in contrast to that seen for several other SR factors, suggesting that C3G is weakly associated with speckles. SC35, and snRNPs differ in their dependence on RNA for retention in speckles (Spector et al., 1991). Proteins in nuclear speckles are known to have specific sequence elements that enable their targeting by binding to RNA and other proteins (Cáceres et al., 1997; Eilbracht and Schmidt-Zachmann, 2001). In the primary sequence of C3G, we could not identify any such features. It is possible that C3G resides in speckles through its ability to interact with other macromolecules present. Since C3G shows properties similar to some snRNPs with respect to extractability, it is possible that localization of C3G to speckles is mediated by snRNPs. Our results show that increased localization to speckles in response to amanitin is dependent on intact cytoskeletal elements. Similar dependence has been shown for other proteins that reside in the speckles (Gieni and Hendzel, 2009). Localization of C3G is also dependent on activity of Clk1 as we observed that enhancing Clk1 activity by its overexpression disrupted the localization of C3G. The possibility of C3G being a substrate of Clk1 is being explored as it is known that Cdk5 phosphorylates C3G (Utreras et al., 2013). Since C3G does not possess an RS motif, it will be interesting to identify interacting partners of C3G that enable it to move in & out of speckles in response to Clk1 mediated phosphorylation.

Rap1, the small GTPase target of C3G localizes to the nucleus, and regulates gene expression (Lafuente et al., 2007), but its association with nuclear speckles was not known. We show that active Rap1 is a component of nuclear speckles, and inactivation of Rap1 dependent signalling alters speckle number, and morphology. These changes appear similar to the effect of transcriptional and splicing inhibitors, suggesting that RNA transcription and/or splicing may be dependent on Rap1 activity.

Rap1 plays a role at cell junctions (Ando et al., 2013) and many junctional proteins and those that regulate actin cytoskeleton have been shown to localize to nuclear speckles (Saitoh et al., 2004). C3G signals to cytoskeleton dependent functions in the cell, and is also known to associate with cytoskeletal elements (Martín-Encabo et al., 2007; Radha et al., 2007). We observed that distribution of C3G, as well as SC35 to speckles is effected upon treatment of cells with Cytochalasin D, a microfilament depolymerizing agent, indicating that enhanced localization of C3G in speckles seen upon inhibition of transcription is dependent on intact cellular microfilaments.

We have earlier shown that inhibition of GSK3β induces import of C3G into the nucleus, where it localizes to non-heterochromatin domains (Shakyawar et al., 2017). We observed that under these conditions, C3G localizes to speckles, suggesting that localization of nuclear C3G to speckles may be dependent on specific physiological stimuli. C3G shows enhanced localization to speckles in differentiated myotubes, and knockdown resulted in reduced intensity of SC35 speckles. The functional consequence of localization to speckles is a role for C3G in regulating splicing activity. While C3G over-expression enhanced splicing activity, knockdown of C3G resulted in reduced splicing. Alternate splicing is an important mechanism that enables myogenic differentiation (Bland et al., 2010). Our earlier work showed that C3G is required for C2C12 differentiation. Therefore, one of the required properties of C3G to enable myogenic differentiation may be its function in alternate splicing. We also observed that cells lacking C3G show reduced presence of SC35 in speckles. C3G could be playing an adaptor function to enable SR proteins like SC35 to home to speckles, and also playing an independent catalytic function in enabling splicing activity. Surprisingly, examination of cell lysates having low levels of C3G due to CRISPR/Cas9 mediated knockdown, showed dramatic decrease in several splicing factors, which could explain reduced splicing activity in these cells. The splicing factors regulated by C3G are ubiquitous components of speckles and essential for constitutive splicing. One likely explanation for reduction in several of these factors, is proteasomal degradation specific to nuclear speckles (Baldin et al., 2008). If C3G functions to inhibit this activity, reduction of several cellular splicing factor protein levels would be expected. An alternate possibility is the ability of C3G to regulate transcription of splicing factors through some presently unknown mechanism. If C3G affects splicing activity in cells, its loss could result in reduced levels of many proteins that are generated from spliced transcripts. These possibilities are being investigated. Though C3G over expression did not alter levels of these factors, it did show significant, but small increase in splicing activity. This could be because C3G is overexpressed only in 30-40 % of cells.

Based on multiple lines of evidence, we describe a novel function for C3G in the nucleus. Localization to splicing factor rich nuclear speckles; dynamic and reversible localization in response to cellular transcription and splicing activity; dependency of localization on intact DNA and RNA; presence of activated Rap1, a target of C3G in speckles; and its requirement for splicing activity enabled us to show that regulation of RNA splicing is an important function of nuclear C3G (Fig. 7E). Ras family GTPases and exchange factors that activate them are involved in multiple signalling pathways regulating cell functions in response to external stimuli. Several of them are known to localize to the nucleus, but their functions in the nucleus are poorly understood. This is the first example of a Ras family GTPase and its exchange factor localizing to, and functioning in nuclear speckles.

## Materials and Methods

Antibodies against C3G(H300), Calnexin, NF-kB, Rap1 and GAPDH were purchased from Santa Cruz biotechnology. Anti-SC35 (S4045-100UL) and Anti-snRNP200 were from Sigma. Anti-SC35 (ab204916), Anti-DHX38 and Anti-Lamin B1 were from Abcam. RNA polymerase-II H5 (Warren et al., 1992) and β-Actin antibodies were from Berkeley Antibody Company and Millipore respectively. U1 snRNP70 (Lerner and Steitz, 1979), U2 snRNP B” (Habets et al., 1985) and Sm snRNP(Y12) (Lerner et al., 1981) antibodies were gifted by David Spector (CSHL, Cold spring Harbour, NY, USA). α-Amanitin, DRB, Cytochalacin D, Nocadazole and LiCl were from Sigma. Leptomycin B was from Santa Cruz Biotechnology. Isoginkgetin was from Merck Millipore. Luciferase reagent was from Promega. Lipofectamine 3000 was from Invitrogen. Horseradish Peroxidase conjugated antibodies were from Amersham GE. Fluorophore conjugated secondary antibodies were from Millipore and Amersham GE.

### Plasmid constructs

Expression vector for Flag tagged human C3G was a kind gift from Dr. Tanaka (Tanaka et al., 1994). SC35-GFP construct was a kind gift from Dr. V. Parnaik (Tripathi and Parnaik, 2008), pTN24 plasmid was gifted by Dr. I.C. Eperon. GFP-Clk1 and GFP-mClk1(K190R) plasmids were gifted by David Spector (CSHL, cold spring Harbour, NY, USA). GFP-RalGDS-RBD and GFP-Rap1GAP constructs were gifted by Dr. P.J. Stork (Carey and Stork, 2002) and Dr. Patrick Casey (Meng and Casey, 2002) respectively.

### Cell Culture, Transfections and Treatments

MDA-MB-231, MCF-7, HEK293T and HeLa cells were cultured in DMEM with 10% FBS, C2C12 cells were maintained in DMEM with 20% FBS under standard conditions and differentiation was carried out as described earlier (Kumar et al., 2015). Lipofectamine 3000 was used for transfections in cells as per manufacturers protocol. Treatments of α-Amanitin, 50 µg/ml, 5 h; LMB, 37 nM, 6 h; Nocodazole, 1µg/ml, 4 h; Cytochalasin D, 0.2 µg/ml, 4 h; IGK 50 µM, 15 h; LiCl, 50 mM, 24 h; and DRB (5,6-Dichlorobenzimadazole-riboside) 25µg/ml, 3 h; were given by incubating exponentially growing cells with respective drug(s) in cell culture medium. Transient heat shock treatment of cells was carried out in an incubator at 42°C for 2 h after addition of pre-warmed medium.

Generation of C3G KO clones in C2C12 cells has been described earlier (Shakyawar et al., 2017). Similar strategy was applied to generate C3G KO clones in MDA-MB-231 cells using commercially available CRISPR/Cas9 KO plasmid (sc-401616) and HDR plasmid (sc- 401616-HDR).

### Immunofluorescence, Image Analysis and Western Blotting

Immunofluorescence was performed as described earlier (Shivakrupa et al., 2003). Co-staining for two antigens was performed by sequential incubation of primary and corresponding secondary antibodies. Parallel coverslips processed without addition of primary antibodies were used as Blank (BL) to show absence of non-specific staining from secondary antibodies. Confocal Z stacks were captured on Leica TCS SP8 Confocal microscope (Leica Microsystems, Germany), and data was analysed using Leica Application Suite and ImageJ software’s. Constant image acquisition parameters were used for capturing images of all samples from a given experiment. All confocal images were captured under 63X or 100X objective of the microscope and were digitally processed for presentation using Adobe Photoshop CS6 software. Quantitation of SC35 positive nuclear speckles was carried out with ImageJ software. Images of cells were imported into ImageJ and a digital brightness threshold was applied to each image. A particle analysis tool available in the ImageJ software was applied to thresholded images and number of structures measuring beyond 0.3 µm in size were obtained as output.

Western blotting was performed as described (Radha et al., 1994). Vilber-Lourmat Chemiluminescence System (Germany) or Carestream XBT autoradiogram sheets were used for detection of ECL signal. ImageJ software was used for quantitation of the western blots and values were normalised to loading control.

### Cell Fractionation and *In situ* extraction

Whole cell (W), Cytoplasmic (C) and Nuclear (N) fractions of cells were prepared as described (Radha et al., 1994). In situ extraction of cells plated on coverslips was performed as described (De Conto et al., 2000) with minor modifications. Cells grown on coverslips were rinsed twice with TM buffer (50 mM Tris-HCl, pH 7.5, 3 mM MgCl_2_), followed by 10 min incubation on ice with TM buffer containing 0.4% Triton X-100, 0.5 mM CuCl_2_, 0.2 mM PMSF and protease inhibitor. Cells were washed and treated with DNase I (40 U/ml) or RNase A (20 µg/ml) for 30 min at 37°C. Cells were subjected to 0.4 M and 2 M NaCl treatment sequentially for 5 min on ice to remove chromatin and RNA. Coverslips were fixed with methanol at each stage and immunostained.

### Splicing activity assay

In vivo splicing activity assays was performed as described (Nasim et al., 2002). pTN24 reporter construct encodes β-gal and luciferase, separated by an intronic sequence (derived from human αs-tropomyosin gene) that contains stop codons. The ratio of luciferase to β-gal reflects the splicing activity in cells. Cells were transfected with the dual reporter plasmid pTN24 in addition to indicated constructs. After 48 hours of transfection, lysates were prepared using reporter lysis buffer and assayed for luciferase activity (Turner designs luminometer TD 20/20), and β-gal assay as per Promega protocols. Luciferase to β-gal ratios were normalised with control to determine relative in vivo splicing activity.

### Statistical analysis

All data is represented as the mean±s.d. Unpaired Student’s t-Test was carried out to obtain the significance of differences in means.

## Acknowledgements

We thank Dr. S. Tanaka, Dr M. Matsuda, Dr. P.J.S. Stork, Dr. Veena K. Parnaik, Dr. I.C. Eperon, and Dr. D.L. Spector for gift of valuable plasmids. We thank N. Rangaraj for expert assistance with confocal microscopy. We thank Kunal Dayma for his critical comments on this manuscript.

## List of abbreviations RapGEF1

RapGEF1: Rap guanine nucleotide exchange factor 1
GSK-3β: Glycogen Synthase Kinase 3 Beta
LMB: Leptomycin B
CRM1: Chromosomal Maintenance 1
IGK: Isoginkgetin
AMA: α-amanitin
DRB: 5, 6-dichloro-1-β-d-ribofuranosylbenzimidazole

